# NeuralTE: an accurate approach for Transposable Element superfamily classification with multi-feature fusion

**DOI:** 10.1101/2024.01.21.576519

**Authors:** Kang Hu, Minghua Xu, Xin Gao, Jianxin Wang

## Abstract

**Motivation:** Classifying Transposable Elements (TEs) at the superfamily level offers deeper insights into species variation and evolution. Recent advancements in third-generation sequencing technologies have made a large number of genomes from non-model species becoming available. However, existing TE classification methods suffer from several limitations, including the necessity to train multiple hierarchical classification models, the incapacity to perform classification at the superfamily level, and deficiencies in both accuracy and robustness. Therefore, there is an urgent need for an accurate TE classification method to improve genome annotation.

**Results:** In this study, we develop NeuralTE, a deep learning method designed to classify transposons at the superfamily level. To achieve accurate TE classification, we identify various structural features of transposons, and use different combinations of k-mers for terminal repeats and internal sequences to uncover distinct patterns. Evaluation on all transposons from Repbase shows that NeuralTE outperforms existing deep learning, machine learning, and homology-based methods in classifying TEs. Testing on the transposons from novel species highlights the superior performance of NeuralTE compared to existing methods. We also conduct TE annotation experiments on rice using different classification tools, and the results show that NeuralTE achieves annotations nearly identical to the gold standard, highlighting its robustness and accuracy in classifying transposons.

**Availability:** NeuralTE is publicly available at https://github.com/CSU-KangHu/NeuralTE.

## Introduction

Transposable Elements (TEs) are mobile DNA sequences within the genome, first identified by Barbara McClintock in her study of genetic instability during maize pigmentation inheritance [1]. These virus-like genomic elements inserted themselves into host genomes millions of years ago in a parasitic manner, which have contributed to the evolution of diverse host traits like internal gestation, memory, coloration, and adaptive immunity [2]. The interplay between TE insertion and host regulation resembles a game of hide-and-seek: TEs evolve to evade host defense mechanisms, while hosts counteract with their mutations, a dynamic central to genome evolution and speciation [3]. Most of the time, TE insertions are detrimental to the host, as their mobility within the genome can cause genetic instabilities, including mutations and chromosome breakage [4]. However, new TE insertions might generate novel functional proteins [5], enhancing the adaptability of mutant individuals to their environment. As TEs accumulate within host genomes, these elements constitute a significant proportion, ranging from 25% to 75% in mammalian genomes. Accurate identification and annotation of transposons are crucial for understanding their roles in genome stability, evolution, and the regulation of gene expression [6].

According to Wicker’s taxonomy [7], TEs are classified into Class I and Class II. Class I elements, also known as retrotransposons, use RNA intermediates for the ‘copy-and-paste’ mechanism during transposition. Based on their structures, retrotransposons can be classified into five orders: Long Terminal Repeat (LTR) Sequences, Dictyostelium Intermediate Repeat Sequences, Penelope-like Elements, Long Interspersed Nuclear Elements (LINEs), and Short Interspersed Nuclear Elements (SINEs) [8]. These orders can be further divided into a total of 16 distinct superfamilies. On the other hand, Class II elements, known as DNA transposons, employ the ‘cut-and-paste’ mechanism without RNA intermediates. Class II elements can be divided into Subclasses 1 and 2. Subclass 1 can be classified into TIR (Terminal Inverted Repeat) and Crypton orders, while Subclass 2 can be classified into the Helitron and Maverick orders. These orders can further be classified into a total of 12 different superfamilies.

Existing tools employ diverse methods for TE classification, such as Hidden Markov Model (HMM) profiles, homology comparison with known TEs, machine learning, and deep learning methods. Nevertheless, the majority of these tools fail to explore the entire hierarchical classification system and cannot classify TEs up to the superfamily level, such as TEclass [9], REPCLASS [10], and PASTEC [11]. TEclass uses 4-mers and 5-mers as features and support-vector machines (SVMs) for TE classification. To classify unknown TEs, PASTEC employs an HMM method, while REPCLASS utilizes a hybrid approach, incorporating homology-based, structurebased, and target site duplication (TSD) methods. There are several classification methods specialize in classifying specific types of transposons. TE-Learner [12] employs the random forest (RF) method to classify TEs, primarily focusing on classifying LTR retro-transposons. LTRdigest [13], LTRclassifier [14], and LTR_retriever [15] are designed to classify LTR elements, while TIR-learner [16] specializes in classifying TIR elements. TEsorter [17] can classify LTR elements into clades, but its performance in classifying other types of transposons remains unsatisfactory.

TE classification was introduced as a hierarchical classification problem [18], and several tools adopt this approach for the classification of unknown TEs, including DeepTE [19] and ClassifyTE [8]. However, training one or multiple classifiers for each level of hierarchical classification system presents challenges in model training and storage. ClassifyTE is a stacking-based approach that combines various machine learning methods for the hierarchical classification of TEs. However, it fails to classify Retrovirus, Penelope, Helitron, and Crypton elements. DeepTE and TERL [20] are both tools using convolutional neural networks (CNNs) for TE classification. The difference between them lies in the fact that DeepTE encodes sequences based on *k*-mer occurrences, while TERL directly encodes the entire TE sequence. RepeatClassifier is a classification module within the TE annotation tool RepeatModeler2 [21]. It performs homology searches against proteins and sequences in known databases to classify TEs, enabling classification up to the superfamily level. However, it relies on known TEs for homology-based classification, making it challenging to classify novel TEs.

In this study, we introduce NeuralTE, a deep learning method designed to classify TEs to the superfamily level. Compared to existing methods, NeuralTE identifies more structural features from TEs, enabling accurate classification of novel transposons. Furthermore, NeuralTE simplifies the classification process by training a single model capable of classifying different types of superfamilies. This eliminates the requirement to train multiple models for different levels of the TE hierarchical classification system, reducing both the complexity of model training and the storage demands. NeuralTE uses two core modules to construct features from TE sequences (**Fig. 1**). Firstly, the sequence feature module encodes TSDs, 5-bp ends of terminal sequences, and TE domain information using one-hot encoding. Secondly, the *k*-mer feature module uses *k*-mer occurrences to construct feature matrices from both terminal repeats and internal sequences of TEs. To discover distinct *k*-mer occurrence patterns in terminal repeats and internal sequences of TEs, different combinations of *k*-mers are used. Subsequently, the features built from these two modules are concatenated and inputted into a three-layered convolutional neural network (CNN) to predict the classification labels of TEs.

**Fig. 1:**
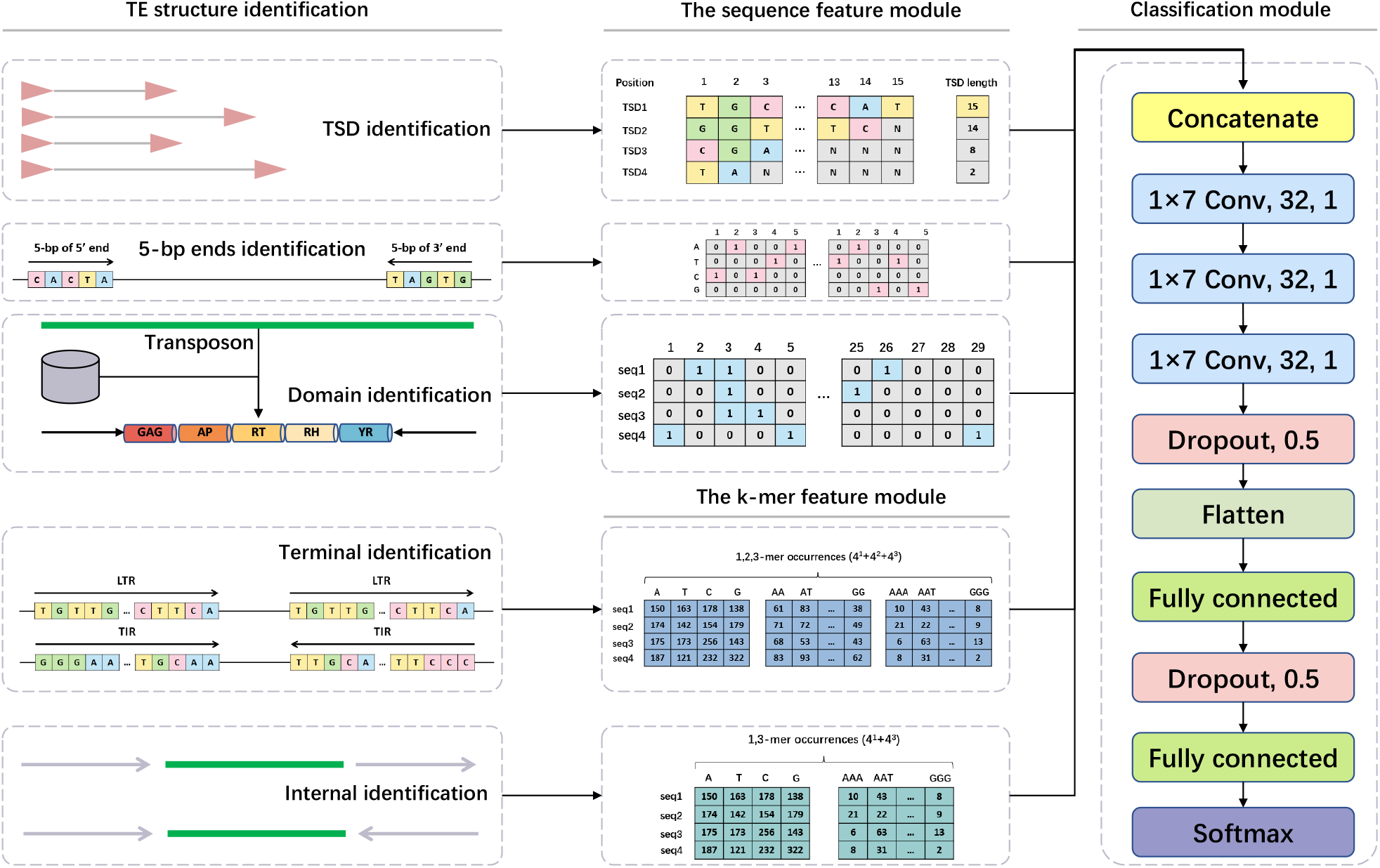
NeuralTE model for TE classification. In the TE structure identification, specific features of transposons are identified, including target site duplications (TSDs), the starting and ending 5-bp sequences, protein domain information, and the terminal repeats and internal sequences. In the sequence feature module, the identified sequence features are converted into one-hot encoding. In the k-mer feature module, the feature matrices are constructed by using k-mer occurrences. In the classification module, (1×7 Conv, 32, 1) denotes a convolutional layer comprising 32 kernels, each with a kernel size of 7, and a stride of 1. In CNN, a kernel refers to a convolution matrix, and the stride represents the movement step size of the convolution sliding matrix at each iteration

We first evaluate NeuralTE on all transposons from Repbase. The results demonstrate that NeuralTE outperforms all existing deep learning and machine learning methods and achieves performance comparable to the state-of-the-art homology-based method, RepeatClassifier. Subsequently, we compare different tools for TE classification in novel species. NeuralTE demonstrates superior performance compared to other tools, outperforming RepeatClassifier in F1-score by 29.55% and 21.64%. Furthermore, when applied to classify transposons in individual species, such as rice, maize, and mouse, NeuralTE demonstrates performance far superior to other tools, aligning with the gold standard for genome annotation. NeuralTE is a user-friendly tool that manages dependencies through *Conda*. All preprocessing steps in NeuralTE are automated, eliminating the need for user intervention.

## Materials and methods

### Datasets

Repbase is a fundamental and well-curated reference database for almost all eukaryotic genome sequence analyses, accessible through a paid subscription [22]. After extensive manual editing and curation, the Repbase database has become reliable enough to serve as the gold standard for testing various TE classifiers. In this study, we downloaded **Repbase** version 28.06 (https://www.girinst.org/repbase/) for model training and testing.

LTR retrotransposons consist of two highly similar long terminal repeats and an internal sequence. Repbase merges two similar long terminal repeats into one to minimize redundancy. To recover fulllength LTRs, we first concatenate the terminal and internal sequences of LTR retrotransposons in Repbase based on sequence headers, resulting in a total of 55,767 sequences from 1,531 species. The hierarchical classification system comprises 28 distinct TE superfamilies. However, since Repbase does not include the Ngaro, VIPER, Maverick, and PiggyBac, we obtain a total of 24 TE superfamilies after converting labels from Repbase to Wicker’s taxonomy [7]. Subsequently, we have acquired 41,198 TE sequences by downloading genome assemblies of 493 species and aligning transposons to these assemblies to identify target site duplications (TSDs) features. Finally, to assess the ability of NeuralTE to identify non-TEs, we have obtained many non-TEs. Considering that NeuralTE performs classification across a total of 24 TE superfamilies, with an average of 1,716 instances per superfamily, we have supplemented the dataset with 1,800 non-TEs, encompassing coding sequences (CDS), random sequences, tandem repeats, satellites, microsatellites, simple repeats, multi-copy genes, and pseudogenes. The inclusion of TEs and non-TEs leads to the formation of **Dataset 1**, consisting of 42,998 sequences derived from 493 species.

### NeuralTE model

As shown in **Fig. 1**, NeuralTE is made up of four modules. The first module identifies specific structures within TE sequences, including TSDs, 5-bp ends of terminal sequences, domain information, terminal repeats, and TE internal sequences. The TE structure information is further fed into feature construction modules. Two modules are tailored to construct distinct features: the sequence feature module constructs features using one-hot encoding from TSDs, 5-bp ends of terminal sequences, and domain information, while the *k*-mer feature module generates features from the sequence of terminal repeats and TE internal sequences through different combinations of *k*-mer occurrences. Subsequently, a CNN-based classification module uses data from both feature modules to predict the TE classification label.

#### TE structure identification The connection of LTRs

In Repbase, LTR elements are split into terminal and internal sequences, such as ‘Copia-62_PHord-LTR’ and ‘Copia-62_PHord-I’. Before identifying TE structures, we concatenate the LTR retrotransposons in Repbase based on the sequence headers to ensure the detection of full-length TEs.

#### The identification of TSDs

We start by downloading an extensive set of genome assemblies from NCBI, including 493 species and containing the majority of sequences in Repbase. To identify TSDs, we conduct BLASTN [23] alignment of TE sequences onto these assemblies, retrieving their copies along with a 20-bp flanking region for each copy. Then, we use regular expressions to search for candidate TSDs within these flanking regions, allowing for a 1-bp mismatch in TSDs longer than 8 bp. To ensure the reliability of identified TSDs, we choose the TSD length that occurs most frequently among all copies. Moreover, given that TSDs occur at the edges of TEs, we designate the positions where TE sequences align with the genome as original boundaries, while boundaries established based on detected TSDs are denoted as identified boundaries. Notably, disparities might arise between original and identified boundaries, with the abbreviated lengths of TSDs potentially resulting in false positives. To ensure the validity of the identified TSDs, TE sequences displaying significant deviations (> 5 bp) between the identified and original boundaries are excluded. These deviations may arise from TE sequences within Repbase being absent at the boundaries or inaccuracies in TSD detection.

#### The usage of 5-bp ends

Different transposons exhibit distinct terminal motifs. For instance, LTR elements commonly present 2-bp palindromic motifs, notably 5’-TG…CA-3’. Helitron elements lack terminal inverted repeats and are inserted into an AT target site, often displaying a canonical terminal structure of 5’-TC…CTRR-3’ (where 5’-TC…CTAG-3’ predominates). In CACTA transposons, we identifies highly conserved CACT(A/G) motifs within their short terminals. Notably, polyA structures are observed at the 3’-ends of 7SL and L1 transposons. Therefore, we use the 5-bp ends of terminal sequences for TE classification.

#### The identification of TE domains

Different transposons exhibit diverse domains, such as different types and orders. For instance, distinguishing between DNA transposons and retrotransposons involves identifying the presence of RT (Reverse Transcriptase) or Tase (Transposase) within the TE sequence. Gypsy and Copia, on the other hand, exhibit variations in the order of RT and INT (Integrase) domains. Therefore, we conduct homology searches between the protein database and TE sequences using BLASTX [24], filtering out bad hits with an evalue cutoff of 1e-20. The protein database containing known TE peptides can be downloaded from RepeatMasker. However, alignment results always present fragmentation, complicating the discovery of complete protein domains within TEs. To resolve this problem, we connect fragmented alignments to establish more continuous domain regions within TE families. Subsequently, we generate a table that maps protein domains to the corresponding TE locations, offering a convenient description of TE protein structures.

#### The identification of terminal repeats

The main difference between LTR (Long Terminal Repeat) and TIR (Terminal Inverted Repeat) elements is that LTRs have long direct repeat sequences spanning from 85 to 5000 base pairs, while TIRs have comparatively shorter terminal inverted repeats, typically ranging from a few base pairs to hundreds of base pairs. To better define the structure of TEs, we use the ‘ltrsearch’ and ‘itrsearch’ tools, included in TE Finder 2.30 (https://github.com/urgi-anagen/TE_finder), to determine whether the TEs exhibit long direct repeats or terminal inverted repeats.

#### The sequence feature module

In the sequence feature module, we set the maximum length of TSDs to 15 bp and use one-hot encoding based on the four ATCG nucleotides. TSDs shorter than 15 bp are padded with zeros. Unknown TSDs or those containing N bases are encoded as all ones. Similar encoding is applied to the 5-bp ends. In NeuralTE, we have defined 29 superfamilies to classify 28 categories from the hierarchical classification system and one ‘Unknown’ category for non-TEs. Since there is a one-to-one correspondence between domains and TE superfamilies, we use vector of length 29 to represent the various types of domains that may occur in a TE sequence, as shown in **Fig. 1**.

#### The *k*-mer feature module

The method of *k*-mer occurrence, used to mining sequence patterns, has been employed in several TE classification approaches [19, 8]. Autonomous LTR and TIR elements have internal sequences that contain the protein domains necessary for their self-replication, typically longer than their terminal sequences. Using different combinations of *k*-mers leads to a richer set of features. Therefore, in contrast to existing methods, we use different combinations of *k*-mers to discover distinct patterns in the terminal and internal sequences of TEs, respectively. In the *k*-mer feature module, we use *k*-mer occurrences to generate feature matrices from the TE sequences (see **Fig. 1**). Let *x*_1_, *x*_2_, …, *x*_*k*_ represents a sequence of *k*-mer (*x*_*k*_ being one of the ATCG nucleotides), the number of features are 4^*k*^.

#### Classification module

In classification module of NeuralTE, we construct a deep CNN model using data from the sequence and *k*-mer feature modules. Our model consists of three convolutional layers with kernel sizes of seven and a stride of one, and does not use any max-pooling layers (**Fig. 1**). Dropout involves randomly deactivating a certain proportion of neurons in a layer during each training iteration, which can help reduce the interdependency between neurons, force the network to learn more robust features and improve its generalization to unseen data.

In NeuralTE, we configure the dropout value to 0.5 after the final convolutional layer. The output from this layer is flattened and served as input to a fully connected layer, where another dropout value of 0.5 is applied. Subsequently, another fully connected layer is used, followed by a softmax output layer to compute probabilities for the classes of input sequences. ReLU function is used in all three hidden layers and the fully connected layer. Model parameters are learned on the training set by minimizing the categorical cross-entropy loss function. We use a batch size of 32, conduct training for 50 epochs, and employ the ADAM optimizer with a learning rate of 0.001.

## Results

### Evaluation of NeuralTE in classifying TEs at the superfamily level

To evaluate the performance of NeuralTE, we have compared our method with five state-of-the-art methods, including two deep learning-based methods TERL [20] and DeepTE [19], two homology-based methods TEsorter [17] and RepeatClassifier [21], and one machine learning-based method ClassifyTE [8] for TE classification.

We randomly select 80% of transposon sequences from **Dataset 1** for model training and the remaining 20% of sequences for testing, and randomly select 20% of transposons from the training set as the validation set. We perform hyperparameter selection based on the validation set. A consistent proportion of transposons from various superfamilies is maintained in both the training, validating, and testing datasets, and all experimental results are based on macro-averaged metrics.

To ensure a fair comparison between NeuralTE and other methods, we have trained models for different approaches on the same training dataset and evaluated them using an identical test set. For example, RepeatClassifier relies on the known library provided by RepeatMasker [25] for TE classification, and we substitute the library in RepeatMasker with our training set to assess its performance. At the same time, since the domain library may contain domains from the transposons to be predicted, there is a potential risk of data leakage during the classification process. Therefore, we filter the domain library to exclude any domain sequences corresponding to species present in the test set. Both the training and prediction phases of the NeuralTE model use the filtered domain library, and the same procedure is applied to RepeatClassifier. An exception is TEsorter, which utilizes its inherent multiple databases for TE classification. As we are unable to replace the built-in database of TEsorter, we consequently assess TEsorter using its default databases and parameters.

DeepTE supports post-processing of prediction results by correcting the classification results through searching for specific domains in transposons. We denote the version of DeepTE using post-processing as DeepTE†. As shown in **Table 1**, NeuralTE outperforms TERL, DeepTE, and TEsorter across all evaluation criteria. TEsorter obtains the lowest F1-score, mainly due to its subpar performance in classifying transposons other than LTR and DIRS. In comparison to TERL, which encodes the entire TE sequence, DeepTE achieves higher performance by using *k*-mer occurrence encoding. DeepTE†, applying TE domains to correct false classifications, achieves a slight improvement compared to DeepTE. RepeatClassifier employs homology-based methods for transposon classification, demonstrating high classification performance on known transposons. Nevertheless, NeuralTE exhibits comparable performance to RepeatClassifier (0.8962 versus 0.9031 for F1-score).

**Table 1:** Evaluation of performance between different TE classification tools on Dataset 1.

| Tools | Precision | Recall | F1-score |
| --- | --- | --- | --- |
| TERL | 0.5201 | 0.4804 | 0.4893 |
| DeepTE | 0.7754 | 0.6268 | 0.6564 |
| DeepTE<sup>1</sup> | 0.7792 | 0.6338 | 0.6624 |
| TEsorter | 0.5591 | 0.3273 | 0.3531 |
| RepeatClassifier | 0.9133 | **0.9022** | **0.9031** |
| NeuralTE | **0.9294** | 0.8754 | 0.8962 |

<sup>1</sup> Applying domains from TEs to correct false classification

Since ClassifyTE fails to classify the Retrovirus, Penelope, Helitron, and Crypton elements, we have filtered out these four types of transposons from **Dataset 1**, resulting in the creation of **Dataset 2**. Subsequently, we perform a random split of the **Dataset 2**, adhering to an 80% training set and 20% testing set principle, and randomly select 20% of transposons from the training set as the validation set. As shown in **Table 2**, NeuralTE demonstrates superior performance compared to ClassifyTE (0.8912 versus 0.7541 for F1-score). At the same time, NeuralTE outperforms ClassifyTE across almost all TE superfamilies. In summary, the results show that NeuralTE has superior performance over TERL, DeepTE, TEsorter, and ClassifyTE, and comparable performance with RepeatClassifier in predicting TE classification at the superfamily level. However, due to RepeatClassifier relying on known TEs for homology-based classification, it faces challenges in accurately classifying novel TEs.

**Table 2:** Evaluation of performance between NeuralTE and ClassifyTE on Dataset 2.

| Tools | Precision | Recall | F1-score |
| --- | --- | --- | --- |
| ClassifyTE | 0.7431 | 0.7892 | 0.7541 |
| NeuralTE | **0.9114** | **0.8773** | **0.8912** |

### Evaluation of NeuralTE on prediction of TEs from novel species

To demonstrate the generalization ability of NeuralTE, we have compared NeuralTE with TERL, DeepTE, TEsorter, and RepeatClassifier in predicting transposons from novel species. We have constructed phylogenetic trees for 493 species from **Dataset 1** to ensure that the species in the test set are novel to the training set. Two different types of test sets are generated based on variations in biological classification and distribution of transposon types. We select flowering plants (57 species) and mammals (113 species) as test sets from the phylogenetic tree, with the remaining species constituting training sets. We select 44 aves species from the training set as the validation set. As shown in **Tables 3 and 4**, there is a significant decline in the performance of all tools when predicting transposons from novel species. Nevertheless, NeuralTE still outperforms all existing methods, including TERL, DeepTE, TEsorter, and RepeatClassifier. Methods based on homology searches, such as RepeatClassifier and TEsoter, often suffer from low recall due to the absence of known transposon in their databases. By incorporating additional features such as TSDs and 5-bp ends, NeuralTE achieves improved classification performance for novel transposons in comparison to existing tools.

**Table 3:** Evaluation of different TE classification tools for predicting transposons in flowering plants (Dataset 3)

| Tools | Precision | Recall | F1-score |
| --- | --- | --- | --- |
| TERL | 0.3659 | 0.3353 | 0.3116 |
| DeepTE | 0.3251 | 0.3205 | 0.2683 |
| DeepTE<sup>1</sup> | 0.3792 | 0.3366 | 0.2966 |
| TEsorter | 0.6051 | 0.3319 | 0.3532 |
| RepeatClassifier | **0.7069** | 0.4792 | 0.4843 |
| NeuralTE | 0.6907 | **0.6855** | **0.6780** |

<sup>1</sup> Applying domains from TEs to correct false classification

**Table 4:** Evaluation of different TE classification tools for predicting transposons in mammals (Dataset 4)

| Tools | Precision | Recall | F1-score |
| --- | --- | --- | --- |
| TERL | 0.2188 | 0.2639 | 0.1713 |
| DeepTE | 0.2545 | 0.3651 | 0.2220 |
| DeepTE<sup>1</sup> | 0.2550 | 0.3672 | 0.2226 |
| TEsorter | 0.2940 | 0.2031 | 0.1209 |
| RepeatClassifier | 0.4985 | 0.5570 | 0.4270 |
| NeuralTE | **0.5421** | **0.6807** | **0.5194** |

<sup>1</sup> Applying domains from TEs to correct false classification

To evaluate the predictive ability of NeuralTE for individual species, we select rice, maize, and mouse for our experiments. We have separated transposons of rice, maize, and mouse within **Dataset 1** for testing, reserving the remaining species for training. We randomly select an individual species from the training set as the validation set. The datasets specific to rice, maize, and mouse are denoted as **Dataset 5, Dataset 6**, and **Dataset 7**, respectively. As shown in **Figs. 2A, B, and C**, NeuralTE outperforms TERL, DeepTE, TEsorter, and RepeatClassifier in both rice, maize, and mouse, achieving F1-scores of 0.7208, 0.9185, and 0.974, respectively. Moreover, NeuralTE demonstrates superior performance across almost all superfamilies compared to other tools. Notably, the Mutator superfamily is recognized as a prevalent contributor to genome instability and genetic variation [26]. NeuralTE achieves a predictive F1-score of 0.94 for Mutator transposons in rice, while RepeatClassifier, TEsorter, DeepTE, and TERL obtain 0.71, 0.32, 0.47, and 0.29, respectively.

**Fig. 2:**
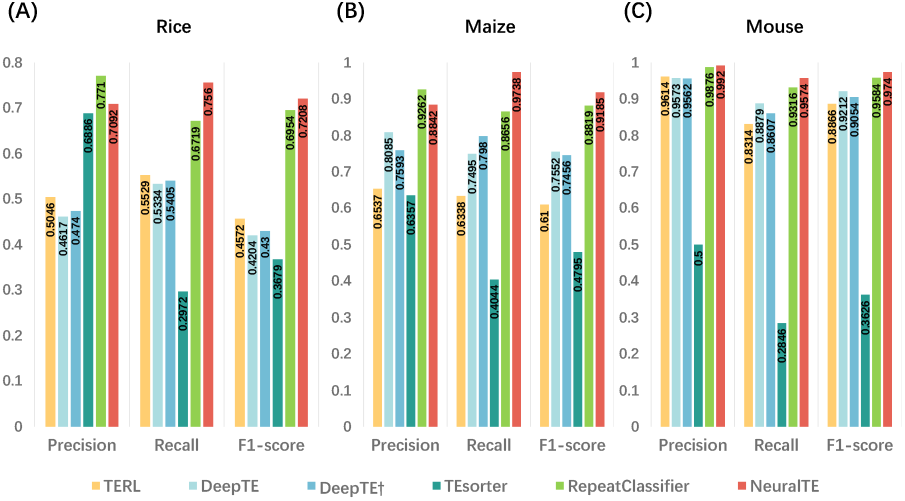
Comparison of the predictive performance of different TE classification tools in rice (**Dataset 5**), maize (**Dataset 6**), and mouse (**Dataset 7**). The predictive performance in rice (**A**), maize (**B**), and mouse (**C**). DeepTE†: Applying domains from TEs to correct false classification

We have compared the number of transposons correctly classified by NeuralTE and RepeatClassifier in both rice, maize, and mouse. As shown in **Fig. 3A**, NeuralTE correctly classifies 2,289 transposons in rice, surpassing RepeatClassifier, which classifies 1,656. Compared to RepeatClassifier, NeuralTE improves by 38.2% in the correct classification of transposons and exhibits a higher overlap with the gold standard. Similar patterns are also observed in maize and mouse. However, due to the lower abundance of TE superfamilies in maize and mouse compared to rice, the gap between NeuralTE and RepeatClassifier narrows (**Figs. 3B and C**).

**Fig. 3:**
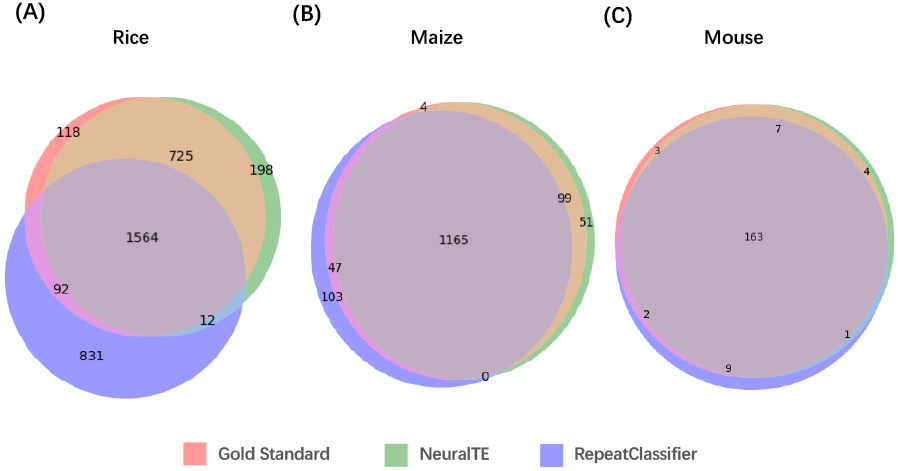
Venn diagram showing the number of transposons correctly classified by NeuralTE and RepeatClassifier in rice (**Dataset 5**), maize (**Dataset 6**), and mouse (**Dataset 7**). The number of transposons in rice (**A**), maize (**B**), and mouse (**C**). The figure indicates that NeuralTE exhibits a higher overlap with the gold standard compared to RepeatClassifier

We have curated eight well-known TIR transposons, including six from rice, one from zebrafish, and one from maize. Each of these transposons has been previously documented in the literature for their significant roles in the functionality and evolution of species. To evaluate the classification performance of different tools on these transposons, we have excluded transposons from rice, zebrafish, and maize species from **Dataset 1**, creating **Dataset 8** as the training set, with these eight transposons serving as the testing set. We randomly select eight TIR transposons from the training set as the validation set. NeuralTE accurately classifies all transposons, whereas other tools exhibit misclassification for certain transposons, notably showing a high error rate for a MITE transposon, mJing. These results show that NeuralTE is more powerful than other tools in predicting the classification of transposons from novel species.

### Using different classification tools for TE annotation

Using different TE classification tools may result in significant differences in TE annotation. To evaluate the genome annotation performance of NeuralTE on novel species, we have compared diverse tools using rice as a model species on **Dataset 5**. As shown in **Fig. 4A**, NeuralTE demonstrates superior predictive performance across almost all superfamilies compared to RepeatClassifier, TEsorter, DeepTE†, and TERL. In the classification of Gypsy and Copia, NeuralTE shows comparable performance with RepeatClassifier. However, NeuralTE exhibits higher performance on other transposons.

**Fig. 4:**
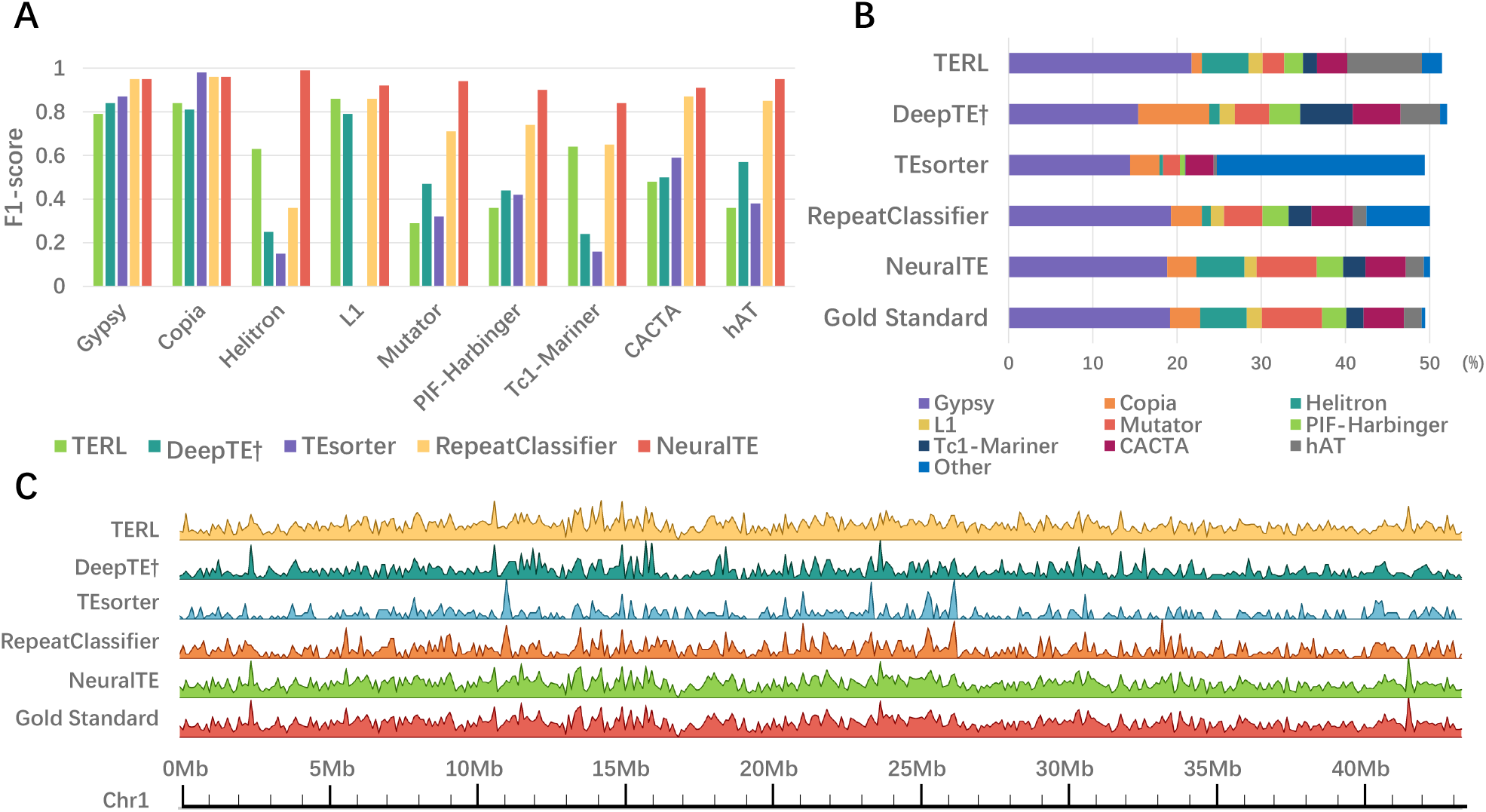
Differential analysis of rice TE classification using various tools (**Dataset 5**). (**A**) Evaluation of the predictive performance of different tools across distinct TE superfamilies using the F1-score metric. TEsorter lacks the ability to predict L1 transposons, resulting in the absence of values. (**B**) Comparison of the percent of genome masked by different superfamilies classified using different tools. (**C**) Comparison of TE density profiles derived from different tools (showcasing Helitron elements in the figure)

Additionally, we use RepeatMasker (version 4.1.1) and libraries classified by different tools to conduct genome annotation using the *-lib* parameter. It is worth noting that we do not use the Dfam library provided by RepeatMasker during the genome annotation process. As shown in **Fig. 4B**, NeuralTE achieves annotation proportions closest to those of the gold standard, Repbase. RepeatClassifier, TEsorter, and DeepTE† show a reduced fraction of the genome masked by Helitron elements. DeepTE† also results in a higher proportion of the genome masked by Copia and Tc1-Mariner elements. TEsorter displays a significant portion of the genome masked by other types of transposons, while TERL indicates a higher fraction of genome masked by Gypsy elements but a lower proportion masked by Copia elements. Furthermore, the transposon density throughout the genome using different tools is displayed. For simplicity, we showcase Helitron density on chromosome 1 of rice. As shown in **Fig. 4C**, NeuralTE achieves a density distribution nearly identical to that of the gold standard. The results show differences in TE annotation when using different classification tools. Notably, NeuralTE outperforms other tools and achieves annotations consistent with the gold standard.

## Discussion

In this study, we develop NeuralTE, a deep learning method to classify transposons into distinct superfamilies. Evaluation of NeuralTE on all transposons from Repbase reveals its superior performance in classifying TEs at the superfamily level compared to existing TE classification methods. To assess the contribution of each feature, we have conducted ablation experiments. All features used by NeuralTE, such as domain, *k*-mer, TSDs, and 5bp-ends, significantly contribute to enhancing model performance. Furthermore, we demonstrate that the integration of HiTE and NeuralTE pipelines enables the analysis of previously unrecorded transposons, potentially augmenting contributions to advancements in plant breeding.

Recent advancements in third-generation sequencing technologies have made a large number of genomes from non-model species becoming available. Our tests on transposons from these novel species demonstrate that NeuralTE outperforms all existing methods in classifying TEs at the superfamily level, showing its remarkable generalization capability. Moreover, when applied to classify transposons in individual species, such as rice, maize, and mouse, NeuralTE demonstrates performance far superior to other tools, aligning with the gold standard for genome annotation.

Existing TE classification methods suffer from several limitations, including the necessity to train multiple hierarchical classification models, the incapacity to perform classification at the superfamily level, and deficiencies in both accuracy and robustness. To solve these problems, NeuralTE makes full use of various TE structure features to enhance the predictive performance of the model. Rather than classifying TEs layer by layer according to the hierarchical classification system, NeuralTE trains a single model to directly classify TEs up to the superfamily level. The results show that NeuralTE achieves higher performance and robustness. In conclusion, NeuralTE demonstrates significant promise as a reliable tool for TE classification. We anticipate that the tool developed in this study will contribute to genome variation research, potentially benefiting applications like crop breeding.

